# The electrostatic allostery could be the trigger for the changes in dynamics for the PDZ domain of PICK1

**DOI:** 10.1101/2020.10.06.328617

**Authors:** Amy O. Stevens, Yi He

**Affiliations:** Department of Chemistry & Chemical Biology, The University of New Mexico, Albuquerque, New Mexico 87131, United States

## Abstract

The PDZ domain is a highly abundant protein-protein interaction domain that exists in many signaling proteins, such as PICK1. Despite the highly conserved structure of the PDZ family, the PDZ family has an extremely low sequence identity, making each PDZ domain unique. PICK1 is the only protein in the human genome that is comprised of a PDZ domain and a BAR domain. PICK1 regulates surface membrane proteins and has been identified as an integral player in drug addiction. Like many PDZ-containing proteins, PICK1 is positively regulated by its PDZ domain and has thus drawn attention to be a potential drug target to curb the effects of substance abuse. The goal of this study is to use all-atom molecular dynamics simulations and the electrostatic analysis program, DelPhi, to better understand the unique interactions and dynamic changes in the PICK1 PDZ domain upon complex formation. Our results demonstrated that the PICK1 PDZ domain shares similar canonical PDZ-ligand hydrogen bonding networks and fluctuations of the carboxylate-binding loop to other PDZ domains. Furthermore, our results are unique to the PICK1 PDZ domain as we reveal that the binding of ligand opens up the binding pocket and, at the same time, reduces the fluctuations of both the central part of the binding pocket and the short loop region between the αA-helix and βC-strand. More importantly, the binding of ligand resulted in charge redistribution at the binding pocket region as well as the N- and C-termini of the PDZ domain that are not a part of the binding pocket. These results suggest that the electrostatic allostery resulted from ligand binding could be the key factor leading to the changes in dynamics which may be associated with the activation of PICK1. Based on these results, an effective drug to target PDZ domain must not only stably bind to the PICK1 PDZ domain but also prevent the electrostatic allostery of the PDZ domain.

## INTRODUCTION

PDZ (PSD-95/Dlg1/ZO-1) domains are highly abundant protein-protein interaction domains found in signaling proteins of various species.^1–3^ The PDZ domain family is one of the largest families of globular domains and have a wide range of functionality in the human proteome. PDZ domains play a critical role in many biological processes such as managing cell polarity, regulating tissue growth and development, trafficking of membrane protein receptors and ion-channels, and regulating cellular pathways.^4–6^ 268 PDZ domains have been identified in 151 unique human proteins.^7^ Despite the vast abundance and broad function of PDZ domains, the secondary structure is highly conserved. The canonical PDZ domains contains six β-strands and two α-helices and has a single binding site in the hydrophobic groove between the αB-helix and the βB-strand, shown in **Fig. 1**. PDZ domains most commonly interact with the last three to five C-terminal residues of target proteins via the carboxylate binding loop that is defined by the highly conserved Gly-Leu-Gly-Phe motif (red).^8^

**Figure 1.**
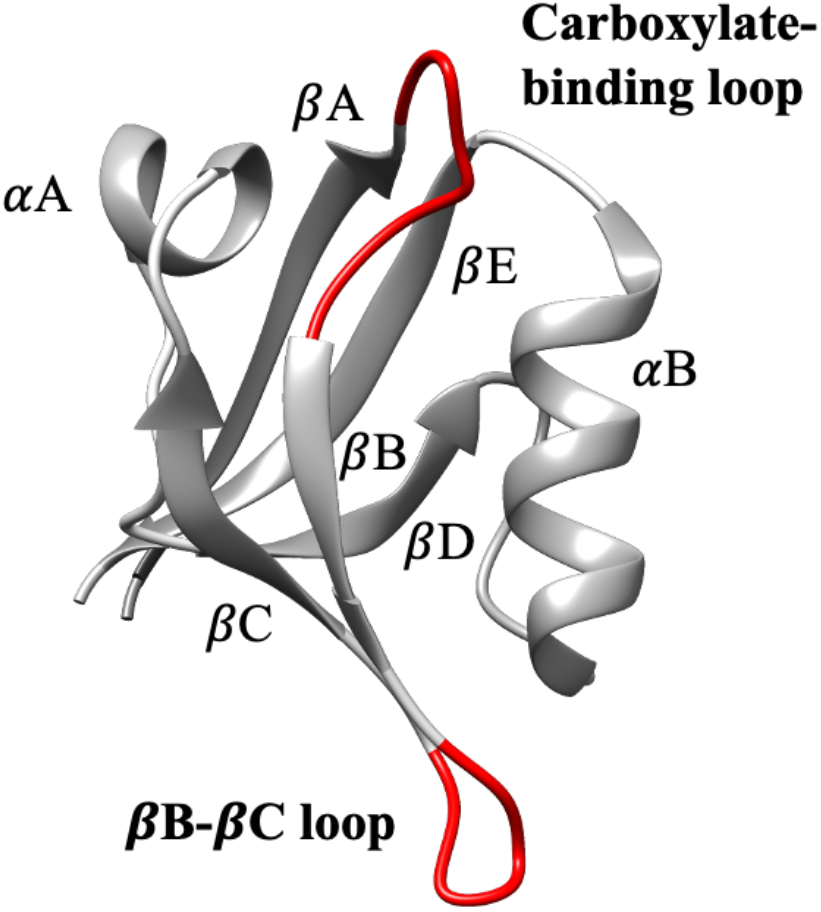
PICK1 PDZ. The hydrophobic binding pocket lies in the groove between the labeled αB-helix and βB-strand. The ligand forms a network hydrogen bonds with the highly conserved carboxylate-binding loop that is shown in red. Additionally, the βB-βC loop is shown in red.

One of many PDZ-containing proteins is Protein Interacting with C Kinase-1 (PICK1). PICK1 is an especially unique PDZ-protein as it is the only protein in the human proteome that is comprised of a both a PDZ domain and a BAR (Bin/amphiphysin/Rvs) domain.^9–11^ PICK1 PDZ takes the canonical binding site in the hydrophobic groove between the *α*B helix and the *β*B strand. The highly conserved carboxylate-binding loop forms a network of hydrogen-bond interactions with the C-terminal tail of its ligands.^8,12^ Functionally, PICK1 is a scaffolding protein that is involved in regulating surface membrane proteins via electrostatic interactions with the cell surface. PICK1 regulates the trafficking of both DAT^13,14^ and AMPA receptor (AMPAR)^15–19^ subunits (GluR2 and GluR3) via interactions with C-terminal residues. The disruption of PICK1-GluR2 interactions abolishes cerebellar and hippocampal long-term depression.^15–17,19,20^ Furthermore, the disruption of PICK1-DAT interactions impairs the behavioral effects of cocaine in mice.^21^ With this, PICK1 has been identified as an integral player in drug addiction.^22^ Consequently, efforts have been made to identify small molecules with the potential to inhibit the function of PICK1.

A deeper understanding of the PDZ domain becomes essential to target PICK1 because PICK1 is activated to perform its function via protein-protein interactions at the PDZ domain. In other words, the biological function of PICK1 is positively regulated by its PDZ domain, and thus the PDZ domain is the target to inhibit the function of PICK1 and reduce the effects of drug addiction.^16,20,23^ Despite novel attempts, successfully developing a PICK1 small-molecule inhibitor with the necessary specificity remains challenging due to the sheer number of unique PDZ domains in the human proteome.^24,25^

Many attempts at targeting PDZ domains for drug development have been made in the past.^26–29^ A recent work by Christensen et al shed light on a drug which can partially reduce the biological function of PICK1.^30^ In parallel, computational groups have aided this effort by performing molecular dynamics (MD) simulations to reveal intradomain interactions of other PDZ domains and PDZ-ligand interactions.^29,31–44^ Furthermore, some efforts have been put towards understanding the structural and energetic allosterism of PDZ domains overall.^31,32,38,39,45,46^ Allosterism refers to the phenomenon that ligand binding at one site results in structural or dynamical changes at a distant site. Computational efforts have highlighted structural fluctuations^31,40,44,46^ and internal redistribution^47^ of energy upon ligand binding. Such proposals have drawn attention as a potential mechanism of global signal transduction across PDZ proteins and opened the possibility of targeting PDZ domains with allosteric drugs. Targeting PDZ domain with allosteric drugs is uniquely challenging as it requires both binding affinity and allosteric effects to be considered simultaneously. ^39, 48–51^

Despite all the efforts given to the PDZ family, few computational efforts have been put towards to the PICK1 PDZ domain. An atomic-level understanding of the key residues involved in allosterism could be the key to assist allosteric drug design.^39^ The purpose of this study is to use a combination of MD simulations and DelPhi^52,53^ to illuminate the interaction mechanisms and atomic-level dynamics between PICK1 PDZ and a representative ligand in the activation of PICK1. Our previous work identified interactions between the component PDZ and BAR domains via replica-exchange molecular dynamics (MREMD) and canonical MD simulations with the coarse-grained UNRES force field.^54^ In this work, we will focus on the changes in structure and dynamics of PDZ domain upon ligand binding. This atomic-level understanding will provide a more complete portrayal of the binding pocket so that PICK1 PDZ can be targeted with the necessary specificity. All-atom MD simulations were performed to model two systems: isolated PICK1 PDZ and the PICK1 PDZ-ligand complex. In this study, the ligand is the final five C-terminal residues (ILVSG) of the carboxyl tail peptide of GluR2. The isolated PICK1 PDZ and the PICK1 PDZ-ligand complex are then compared to illuminate the atomic-level dynamic changes in PICK1 PDZ upon ligand binding. We provide a thorough analysis of both dynamic changes and electrostatic allosterism of the PICK1 PDZ domain. This work contributes a more complete understanding of allosterism across the PDZ domain.

## METHODS

To compare the dynamical changes in PDZ upon the binding of ligand, two sets of simulations were performed. One set includes three trajectories simulating the free PDZ without the presence of ligand. In parallel, the second set also include three trajectories simulating the PDZ-Ligand complex. The experimentally determined complex structure of PDZ (PDB ID: 2PKU) was used to generate the starting structures for all simulations. The most recently developed CHARMM36m^55^ force field with explicit solvents was used in all simulations with the Groningen Machine for Chemical Simulations (GROMACS) package^56–58^, version 2018.3. Counter ions (Na^+^ or Cl^-^) was added to neutralize the whole system at 300 K. All systems were prepared using CHARMM-GUI^59,60^. The steepest-descent minimization and a 1-ns MD equilibrium simulation were carried out to generate equilibrated starting structures for the MD simulations. All bonds with hydrogen atoms are converted to constraints with the algorithm LINear Constraint Solver (LINCS) ^61^. A Nose–Hoover temperature thermostat^62,63^ was used in all simulations. The time step was 2 fs, and snapshots were taken every 100 ps. 110Å cubic water box was used and run for a total of 700 ns per trajectory, a total 2.1*µ*s (3 × 0.7*µ*s) per system.

The electrostatic potential was calculated by DelPhi^52,53^. The charge and the radius of the atoms are assigned using the program pdb2pqr^64^, with the force field CHARMM27^65^. The electrostatic potential distribution of the protein was then generated using Delphi. For Delphi calculation, parameters were used as suggested in previous work^66,67^. For examples, the dielectric constants for protein and water were set to be 2 and 80, salt concertation: 0.15 mM, resolution: 2 grids/Å, probe radius: 1.4 Å and protein filling the DelPhi calculation box (perfil): 70. Visualization of the electrostatic potential distribution was performed in Chimera^68^ with color scale from −1 to 1 kT/Å.

## RESULTS

The isolated PICK1 PDZ and the PICK1 PDZ-ligand complex were compared to better understand the dynamic changes resulting from ligand binding. DSSP^69,70^ analysis was used to investigate the stability of the helical regions of each system. Based off the starting structures, isolated PICK1 PDZ and the PICK1 PDZ-ligand complex both contain alpha helices, *α*A and *α*B, that span residues Thr56-Gly62 and residues Thr82-Val93, respectively. However, the stability of each helix is unique. In our simulations, the frequency of stable *α*A and *α*B helices was ∼75% and 100%, respectively. It is clear that *α*A is less stable than *α*B. Moreover, complex formation does not appear to affect the presence of the *α*-helices. These results are in agreement with previously determined crystal structures of PDZ domains.^3^

Residue-level interactions were analyzed for the combined trajectories of both the isolated PICK1 PDZ and PICK1 PDZ-ligand complex. A contact is formed if the distance between any two heavy atoms of any two residues, which are more than five residues apart in the sequence, is under 0.55 nm. To compare the stability of each contact, contacts are calculated for each frame of the combined trajectories and then summed to obtain the frequency such contact appears in the combined trajectories. Fig. 2A and 2B displays the contact maps of PICK1 PDZ in the isolated and complex states, respectively. Most notably, intradomain contact decreases at the carboxylate-binding loop and *β*B strand (residues Leu32-Gly39) upon ligand binding. Fig. 2C displays the contact map between PICK1 PDZ and the ligand in complex. The contact at the carboxylate-binding loop and *β*B strand that was lost within the isolated PICK1 PDZ is recovered by the ligand in the complex. Fig. 2D displays a representative frame of the conformational change of the *β*B strand upon complex formation via an overlay of the isolated PDZ domain (blue) and PDZ domain in complex with ligand (red). Upon ligand binding, the *β*B-strand shifts away from the *α*B helix as contact dissipates. The ligand disrupts typical protein interactions to form canonical interactions with the carboxylate-binding loop.

**Figure 2.**
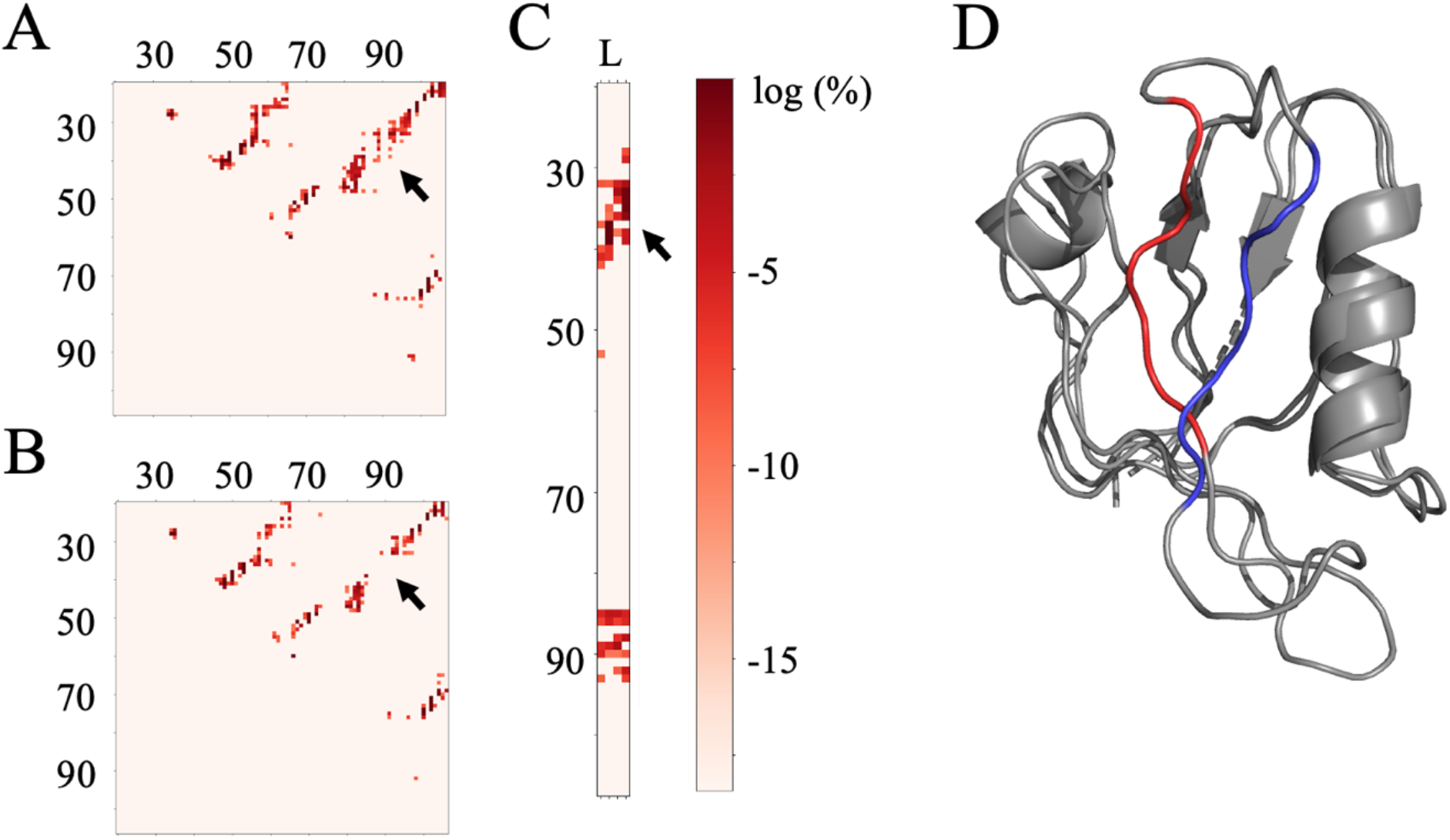
Contact maps. (A) Isolated PICK1 PDZ intradomain contacts. (B) PICK1 PDZ complex intradomain contacts. (C) PICK1 PDZ-ligand (L) contacts. The ligand binding disrupts intradomain interactions at the carboxylate-binding loop as indicated by the arrows in A, B and C. Note, all the contacts are normalized by the total number of the frames in the combined trajectories. (D) Representative overlay of isolated PICK1 PDZ domain (blue) and the PICK1 PDZ-ligand complex (red).

A histogram of the distance separation between the center of *β*B strand (C*α*of residue Ser36) and the center of *α*B helix (C*α*of residue Ala87) reveals the conformational changes at the binding pocket upon complex formation, as shown in Fig. 3. The distance separation between the *β*B strand and the *α*B helix of the isolated PICK1 PDZ domain (light gray) has a wider variance compared to the PICK1 PDZ-ligand complex (red). However, ligand binding appears to result in a more open binding pocket because of the need to fit the large ligand. This can be seen from the shift of the peak of the distance distribution. It should be noted that opposing results were observed in other PDZ domains.^31^ Lu et al suggested that the *β*B strand and the *α*B helix of the PSD-95 PDZ2 domain to shift towards from the binding pocket and thus reduce distance separation upon complex formation.^31^

**Figure 3.**
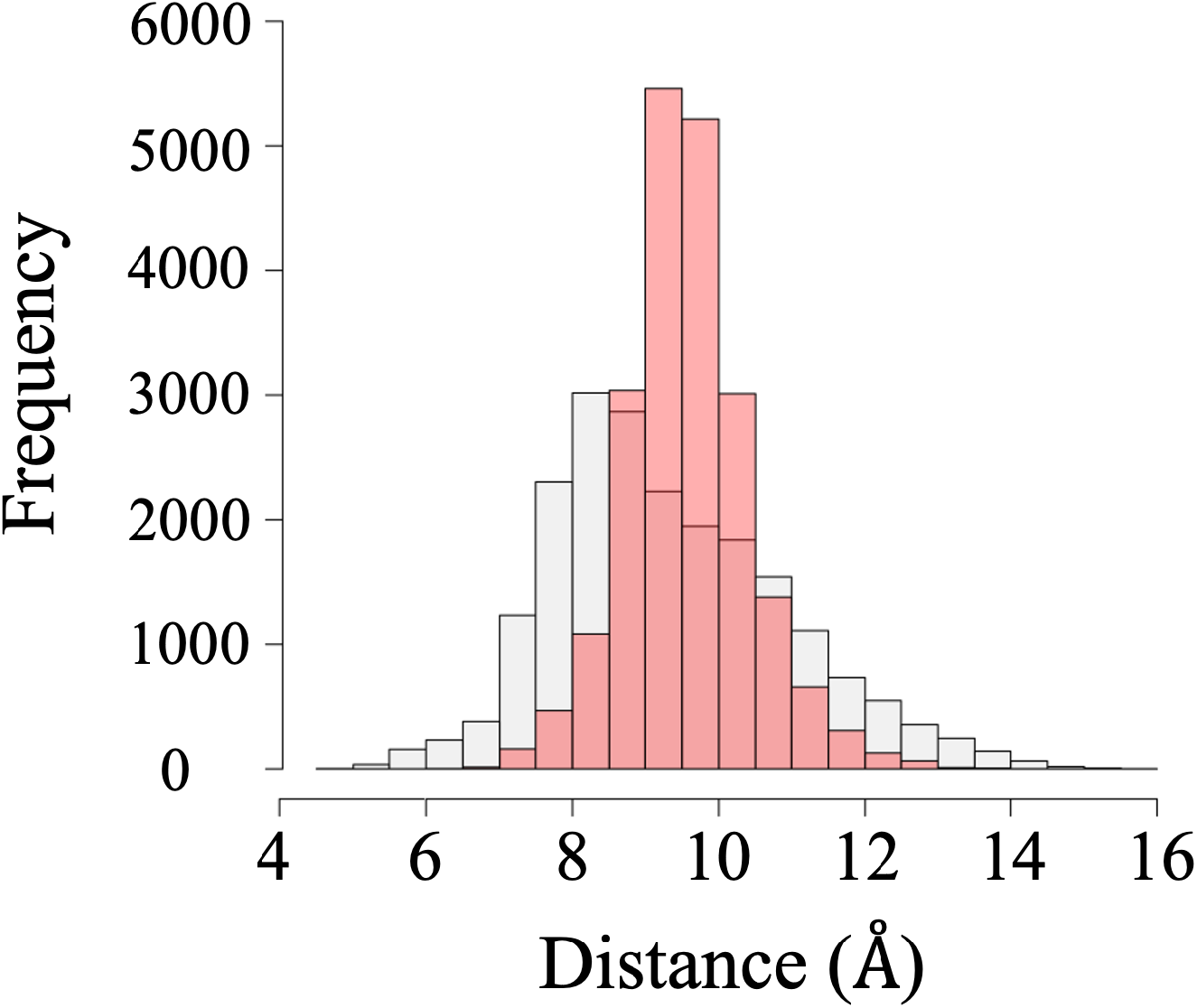
Histogram of distance separation between the βB strand and the αB helix that compose the binding pocket. The light gray bars represent the isolated PICK1 PDZ domain. The red bars represent the PICK1 PDZ-ligand complex.

Hydrogen bond analysis at the binding pocket was performed to determine interactions with the ligand. Our results show the canonical PDZ-ligand hydrogen bond interactions (Fig. 4). The hydrogen bonding network calculated from our simulations were translated to Fig. 4C which display hydrogen bonding interactions in the complex state in two dimensions. Our system displays canonical Class II PDZ interactions.^4,71^ More interestingly Ile37 (PDZ) and Val123 (ligand) form a much stronger pair of donor-acceptor hydrogen bonds than any other hydrogen bond in the binding pocket, as shown in Fig. 4A and 4B. This pair of interactions is essential to the stability of complex formation.

**Figure 4.**
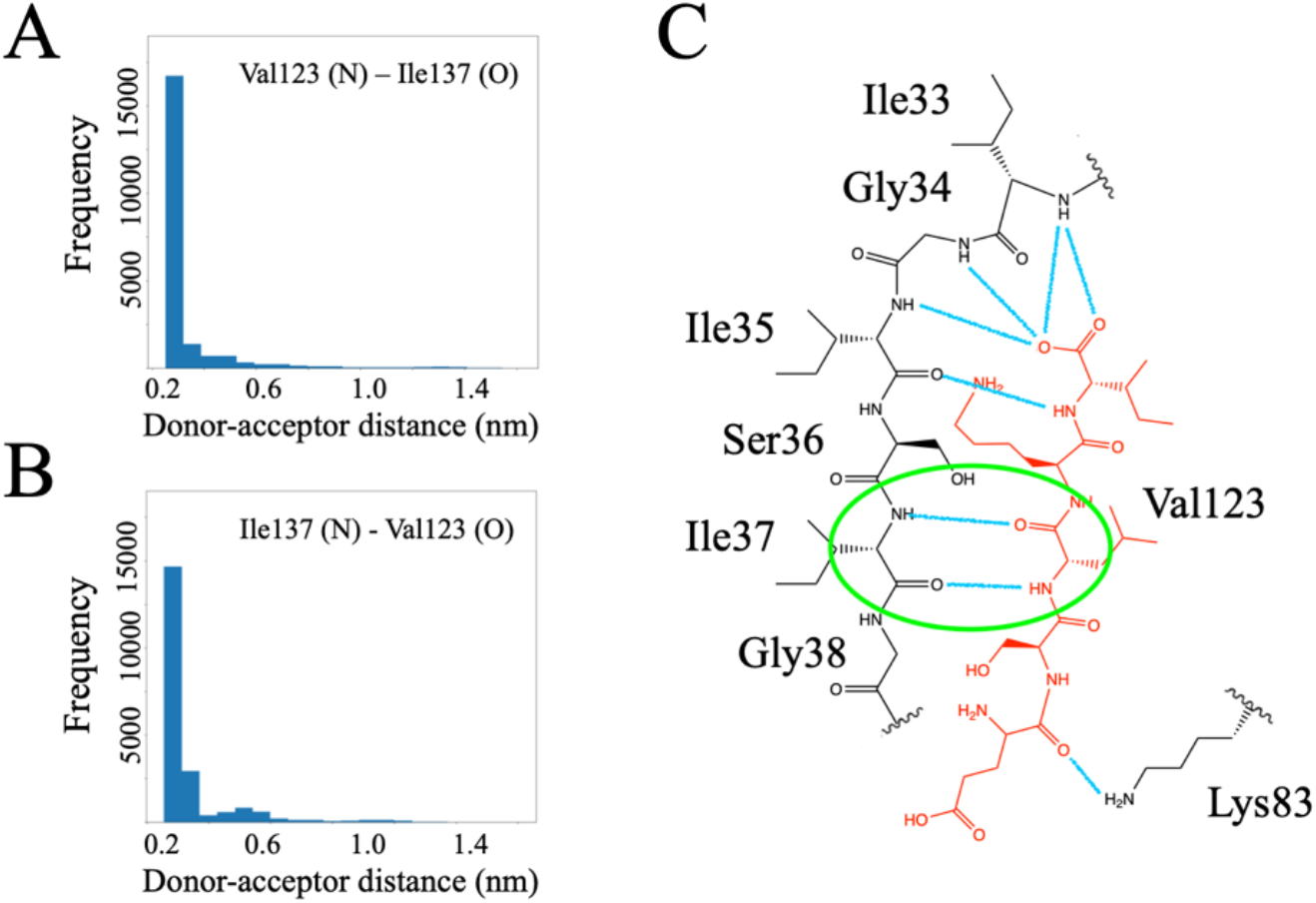
Hydrogen bonds in PICK1 PDZ-ligand complex.

Principal component analysis (PCA) and RMSF were used to explore the differences in movement between the two systems. PCA was performed in efforts to explore changes in the motion of PICK1 PDZ upon ligand binding. Major movements occurred at the carboxylate-binding loop, the *α*A helix and the *β*B-*β*C loop, as shown below in Fig. 5. The major movements occurring at the *α*A helix decrease in the PICK1 PDZ-ligand complex. It should be noted that similar structural allosterism has also been identified at the *α*A helix upon ligand binding in other PDZ domains.^31,46^ Additionally, the major movements of the *β*B-*β*C loop differ between isolated PICK1 PDZ and the PICK1 PDZ-ligand complex. In isolated PICK1 PDZ (Fig. 5A), residues Pro57-Gly62 form the *β*B-*β*C loop that closes towards the binding pocket. Oppositely, in the PICK1 PDZ-ligand complex (Fig. 5B), this loop opens away from the binding pocket. The flexibility of the carboxylate-binding loop and the *β*B-*β*C loop in the complex state may result in the weakening of hydrogen bonds in these regions of the binding pocket. The key pair of donor-acceptor hydrogen-bonds between Ile37 (PDZ) and Val123 (ligand) may be able to form a strong interaction due to the reduced fluctuations of the *β*B-strand at this position.

**Figure 5.**
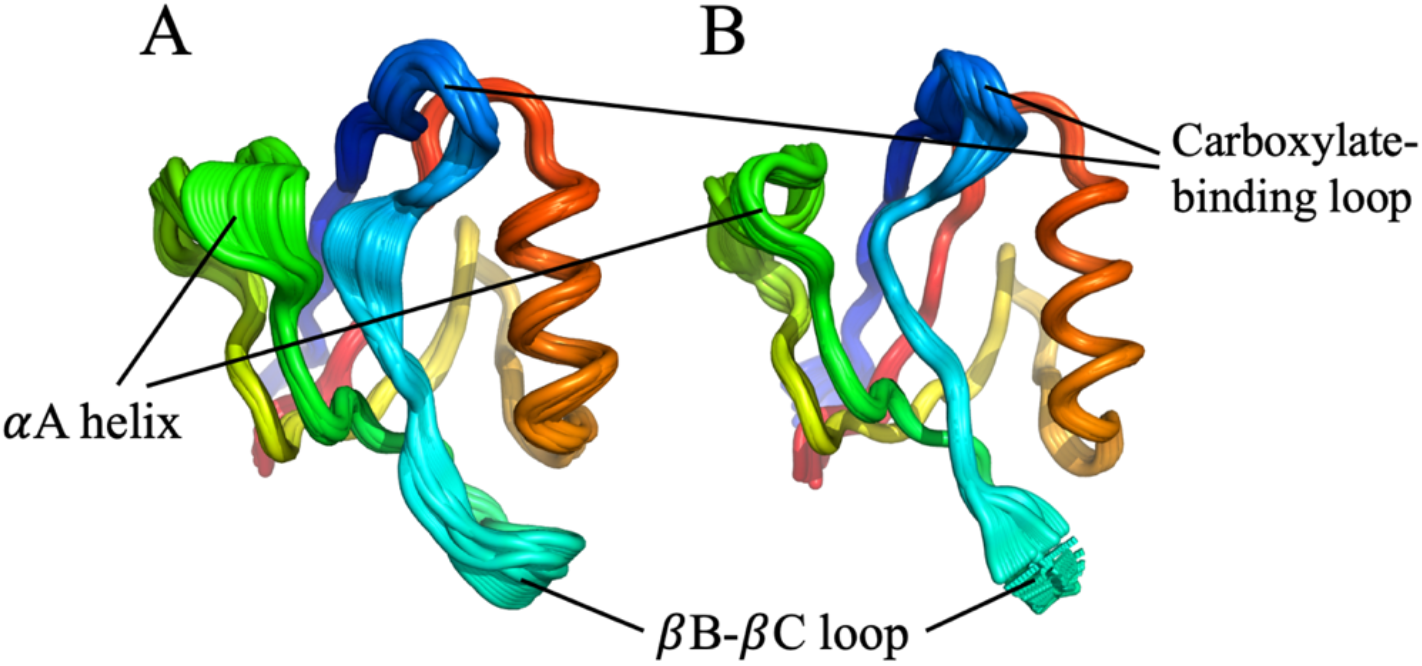
Regions with major changes movements in isolated PICK1 PDZ (A) and the PICK1 PDZ-ligand complex (B). Upon ligand binding, the carboxylate-binding loop (royal blue) closes toward the binding pocket upon ligand binding, the αA helix decreases fluctuations, and the βB-βC loop opens away from the binding pocket.

RMSF analysis was performed to further explore differences in motion between the two systems. The cross-correlation map is an effective tool to identify regions of interaction that are affected by complex formation. The isolated PDZ domain (Fig. 6C) has region of positive coordination (residues Gly34-Ile35 with residues Asp54-Thr56) that is absent in the PICK1 PDZ-ligand complex (Fig. 6A). Additionally, the isolated PDZ domain (Fig. 6C) has a region of negative coordination (residues Thr24-Gln26 with residues Asp54-Thr56) that is reduced in the PICK1 PDZ-ligand complex (Fig. 6A). Furthermore, as shown in Fig. 6B, various regions of residues have a reduced fluctuation in the PICK1 PDZ-ligand complex (red dashed line in Fig. 6B) than in isolated PICK1 PDZ (blue solid line in Fig. 6B). The formation of the complex reduces the fluctuation of the *β*B strand, *α*A helix, and *β*B helix or residues Ile35-Gly38, Thr56-Gly62, and Thr82-Val93, respectively. Notably, the RMSF plot shows a decrease in fluctuation from the isolation PICK1 PDZ to the PICK1 PDZ-ligand complex at the *β*B-*β*C loop. These results agree with our PCA observations. Overall, our results reveal an increased flexibility of the PICK1 PDZ domain in the absence of ligand. This echoes similar results that have been overserved in other PDZ domains.^46^

**Figure 6.**
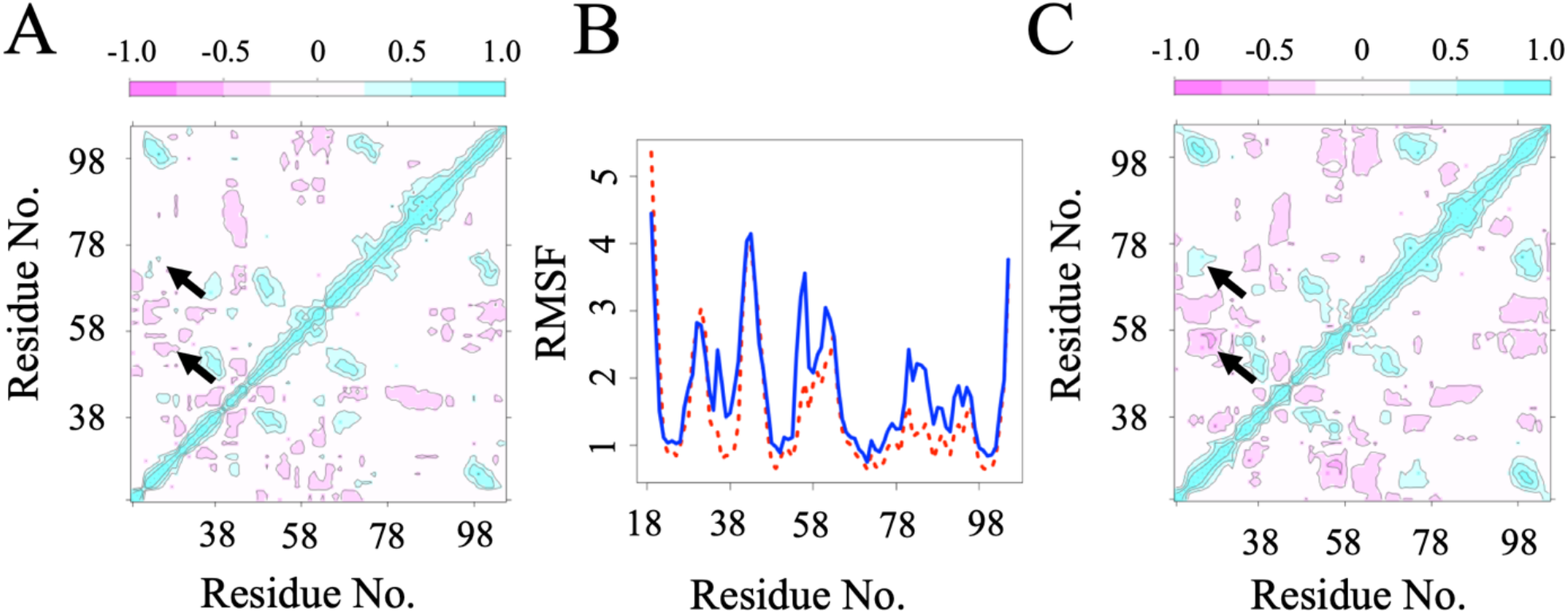
RMSF and cross-correlation maps. (A) Cross-correlation map for PICK1 PDZ-ligand complex. (B) RMSF comparing isolated PICK1 PDZ (solid blue) and PICK1 PDZ-ligand complex (dashed red). (C) Cross-correlation map for isolated PICK1 PDZ.

Functionally, PICK1 is involved in regulating surface membrane proteins via positively charged residues of the BAR domain interacting with the negatively charged lipid bilayer ^72,73^. However, it is difficult to track charge distribution changes, especially when the surrounding solution and metal ions must be considered. In efforts to understand the PDZ domain’s role of activation in this surface-potential driven process, we used DelPhi to analyze the changes in surface potential of PICK1 PDZ upon ligand binding, shown in Fig. 7. Fig. 7A displays a cartoon view of the PICK1 PDZ domain with its binding ligand in spheric view. DelPhi can model the electrostatic potential of biological macromolecules with the presence of water and ionic strength even when proteins have irregular shapes. The green circles highlight the locations where the major charge redistribution occurs. Fig. 7B shows the surface potential of isolated PICK1 PDZ, and Fig. 7C displays the surface potential of the PICK1 PDZ-ligand complex. Major changes in charge distribution are seen both at the binding pocket and at the N- and C-termini of the ligand. Similar results were observed for other PDZ domains.^47^ It should be restated that identical structures for the PICK1 PDZ domain was used in each panel in Fig.7. Panel C differs from panel B only by the presence of the ligand. Sensibly, the ligand induces electrostatic changes at the binding pocket, but it also results in a propagation of charge redistribution to induce electrostatic changes at the N- and C-termini that are distant from the binding pocket. The PICK1 PDZ domain adheres the interdomain linker and BAR domain via its C-terminal to construct the complete PICK1 protein. The changes in surface potential distribution upon ligand binding that occur near the C-terminal of the PDZ domain have the potential to affect its dynamics as well as its subsequent PDZ-BAR interactions.

**Figure 7.**
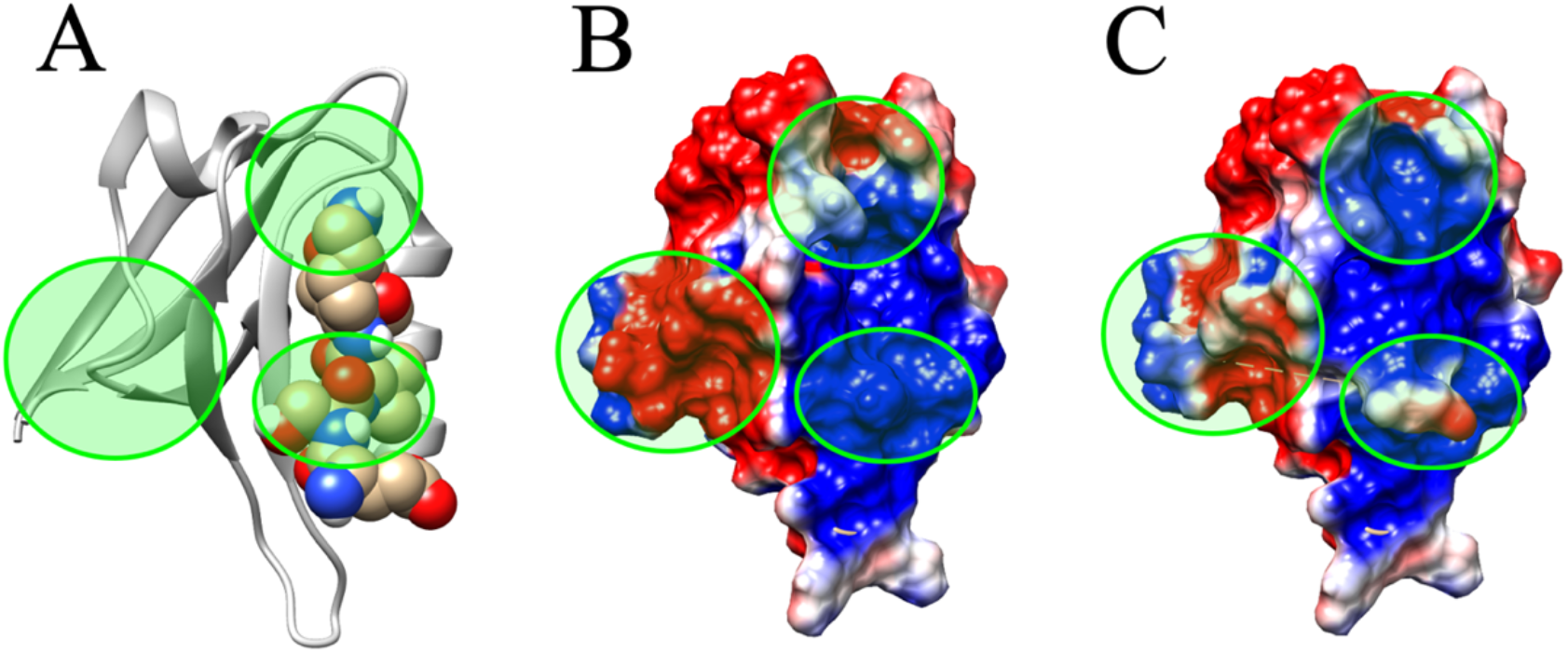
Electrostatic surface potential. (A) PICK1 PDZ (shown in cartoon) complex with ligand (shown in sphere). (B) Surface potential of isolated PICK1 PDZ. (C) Surface potential of the PICK1 PDZ-ligand complex. Green circles highlight regions of major charge redistribution. The positive and negative residues are colored in blue and red, respectively

The PSD-95 PDZ domain is a widely studied PDZ domain. It has a very similar tertiary structure (RMSD 3.178 Å) but shares a 24.4% sequence identify with PICK1 PDZ domain. Structural analysis was performed against PSD-95 PDZ to understand the uniqueness of PICK1 PDZ. Fig. 8A uses normal mode analysis to reveal differences in fluctuation. It should be noted that such fluctuation patterns have been also observed for other PDZ domains.^31^ The carboxylate-binding loop (peak 1 in Fig. 8A) and the *β*B-*β*C loop (peak 2 in Fig. 8A) greatly differ between PSD-95 PDZ and PICK1 PDZ. Fig. 8B shows an overlay of PSD-95 PDZ (blue) with PICK1 PDZ (red). The secondary structures of each domain are in good agreement, but structural differences arise at the loop regions. The normal mode analysis suggests larger fluctuations for the PICK1 PDZ domain compared to the PSD-95 domain, in particular, at the two biological-function related loops.

**Figure 8.**
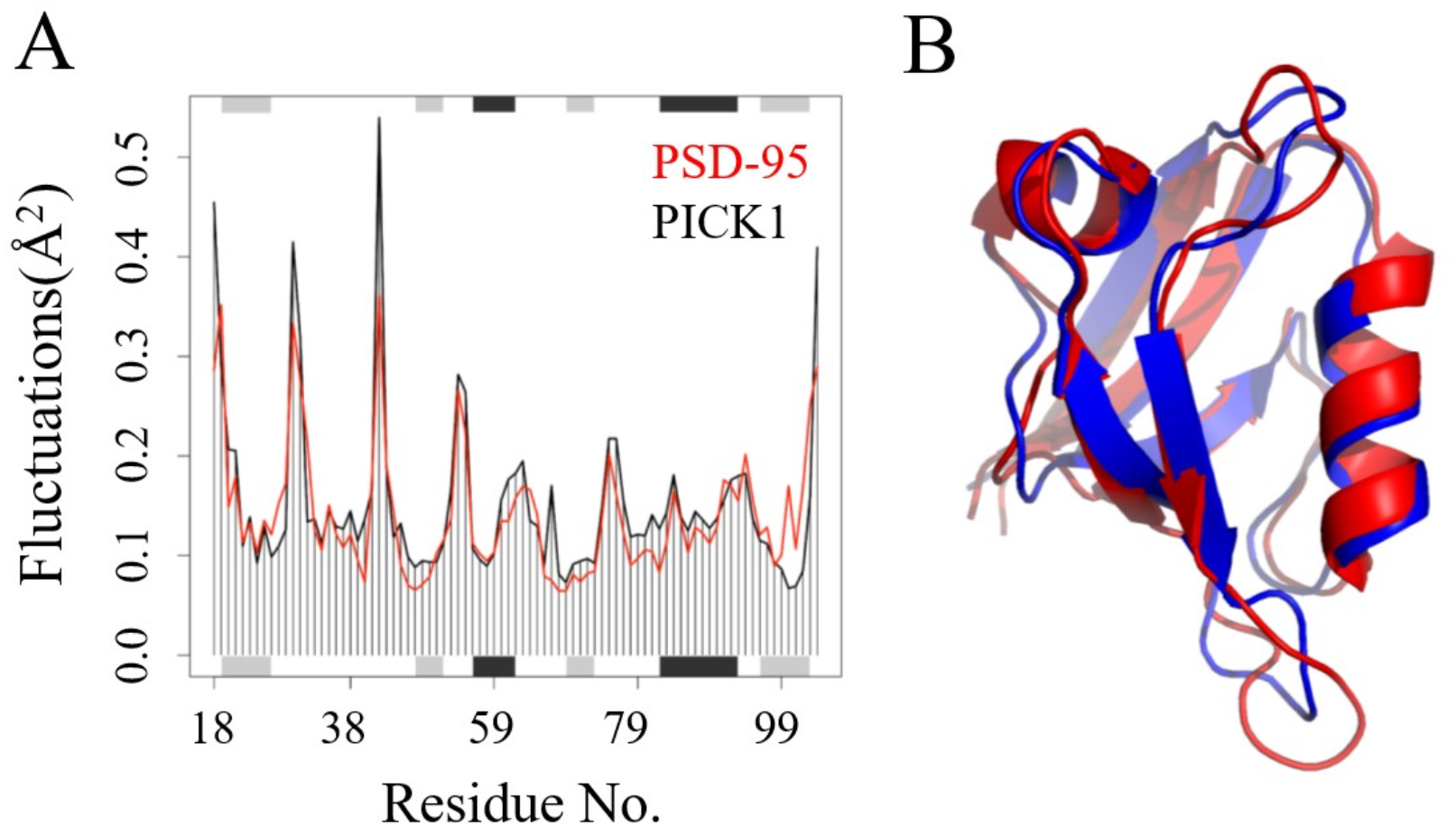
Comparison of PSD-95 PDZ and PICK1 PDZ. (A) Normal mode analysis of PSD-95 PDZ (red) and PICK1 PDZ (black). (B) Overlay of PSD-95 PDZ (blue) and PICK1 PDZ (red).

## DISCUSSION

The purpose of this work is to provide a complete picture of both dynamic changes and electrostatic allosterism of the PICK1 PDZ domain upon complex formation with ligand. We found that (1) ligand binding reduces fluctuations of the *β*B strand and thus allows for stronger hydrogen bonding interactions in the binding pocket, (2) binding of ligand can lead to a wider opening of the binding pocket, and (3) ligand binding results in the redistribution of charge about the PDZ domain which can lead to changes in dynamics of the PDZ domain. Our results provide insight to the structural and electrostatic changes that are induced via complex formation. Ultimately, these results may support an understanding of how ligand binding works to activate PICK1 to perform its biological function.

Canonical Class II PDZ-ligand interactions with the carboxylate-binding loop were seen in the hydrogen bond analysis. While these basic results are in good agreement with previous literature, our analysis went further to reveal unique differences in the strengths of the interactions that compose the hydrogen bonding network. The Ile37-Val123 hydrogen-bond pair has a much greater strength than other interactions in the network because of its unique position in the binding pocket. Here, it seems that the reduced fluctuations of the *β*B strand upon ligand binding allows for increased stability of Ile37-Val123 interactions. With the eventual goal to identify a small-molecule inhibitor in mind, this pair of hydrogen bonds may be essential in drug development. Second, principal component analysis revealed the carboxylate binding loop and the *β*B-*β*C loop open and close toward the binding pocket upon complex formation. Previous literature has confirmed the importance of the carboxylate-binding loop in complex formation and revealed structural allosterism in *β*B-*β*C loop.^31,44,46^ Additionally, previous literature reports that the less studied, poorly conserved *β*B-*β*C loop may have a role in ligand binding.^41,74^ We suspect that the unique conformational changes of the *β*B-*β*C loop as well as its unique structural differences against other PDZ domains may prove valuable in the development of small-molecule inhibitors.

A major challenge in targeting PDZ domains lies in the specificity of small-molecule inhibitors. The abundance of PDZ domains in signaling proteins in the human body requires that small-molecule inhibitors must have the selectivity to target a specific PDZ domain in a distinct signaling protein. A recent work reports a small molecule inhibitor, Tat-P_4_-(C5)_2_, with both strong affinity and specificity to the PICK1 PDZ domain.^30^ Furthermore, Tat-P_4_-(C5)_2_ demonstrates efficacy in reducing addiction-like behavior in mice.^75^ Tat-P_4_-(C5)_2_ interacts with the binding pocket of the PDZ domain as well as negatively charged residues on of *β*B-*β*C loop which introduces an additional layer of stabilization. Our structural analysis against PSD-95 PDZ highlights the unique structure of PICK1 PDZ at the *β*B-*β*C loop. Furthermore, our PCA reveals structural changes at the *β*B-*β*C loop upon complex formation. The uniqueness of PICK1 PDZ at the *β*B-*β*C loop may allow Tat-P_4_-(C5)_2_ to have strong specificity forwards PICK1 PDZ as the inhibitor targets this region of the protein.

PCA and electrostatic surface potentials display ligand-induced allosteric effects across PICK1 PDZ. These allosteric effects may hold the key to understanding how the PICK1 PDZ activates PICK1 to perform its function. Previous literature reports ligand-induced allosteric effects in other PDZ domains.^4,8,28,47,76–81^ Individual PDZ domains have displayed both conformational allostery^39^ and electrostatic allostery.^47^ Our results reveal that the PICK1 PDZ domain simultaneously presents both conformational and electrostatic allostery upon complex formations. Ligand binding results in reduced fluctuations of the *α*A helix, reduced fluctuations in the carboxylate-binding loop, altered structural orientation of the *β*B-*β*C loop, and, perhaps most notably, redistribution of charge across the surface of the N and C-termini. Structurally, our results are in agreement with previous studies revealing allosterism in PDZ domains. The *α*A helix, carboxylate-binding loop, and the *β*B-*β*C loop have each been identified as regions of the PDZ domain structurally affected by allosterism upon ligand binding.

One previous study considered the allostery of the PSD-95 PDZ3 domain by quantifying the electrostatic interaction energy at each residue.^47^ The group proposed that the allosteric structural changes upon ligand binding are a result of an internal redistribution of electrostatic charge upon complex formation. Furthermore, the work suggested allosteric modulation of the PSD-95 PDZ3 domain may propagate electrostatic interactions and lead to a global response from the entire PSD-95 protein. Here, we demonstrate that ligand binding in PICK1 PDZ may trigger this global response. We reveal the inherent electrostatic allostery of the PICK1 PDZ domain upon complex formation. Our results reveal the redistribution of charge at the N- and C-termini, suggesting this charge redistribution may affect tethered modular domains. PICK1 performs its biological function of regulating surface membrane proteins by interacting with the negatively charged lipid bilayer of the cell and is positively regulated by its PDZ domain. The redistribution of charge at the C-terminal of the PDZ domain that our study revealed may then propagate through the linker and BAR domain and ultimately result in charge-charge interactions at the surface of the lipid bilayer as PICK1 performs its function. We hope to search deeper into this possibility by studying the dynamic changes in the PICK1 linker and BAR domain upon complex formation.

## ACKNOWLEDGEMENTS

This work was supported by the Substance Use Disorders Grand Challenge Pilot Research Award, the Research Allocations Committee (RAC) Award, and the startup fund from the University of New Mexico. Molecular graphics and analyses performed with UCSF Chimera, developed by the Resource for Biocomputing, Visualization, and Informatics at the University of California, San Francisco, with support from NIH P41-GM103311.

